# Promoter CpG density predicts downstream gene loss-of-function intolerance

**DOI:** 10.1101/2020.02.15.936351

**Authors:** Leandros Boukas, Hans T. Bjornsson, Kasper D. Hansen

**Affiliations:** Human Genetics Training Program, Johns Hopkins University School of Medicine; McKusick-Nathans Department of Genetic Medicine, Johns Hopkins University School of Medicine; Department of Pediatrics, Johns Hopkins University School of Medicine; Faculty of Medicine, University of Iceland; Landspitali University Hospital; Department of Biostatistics, Johns Hopkins Bloomberg School of Public Health

**Author notes:** Correspondence to (KDH), (HTB).

## Abstract

The aggregation and joint analysis of large numbers of exome sequences has recently made it possible to derive estimates of intolerance to loss-of-function (LoF) variation for human genes. Here, we demonstrate strong and widespread coupling between genic LoF-intolerance and promoter CpG density across the human genome. Genes downstream of the most CpG-rich promoters (top 10% CpG density) have a 67.2% probability of being highly LoF-intolerant, using the LOEUF metric from gnomAD. This is in contrast to 7.4% of genes downstream of the most CpG-poor (bottom 10% CpG density) promoters. Combining promoter CpG density with exonic and promoter conservation explains 33.4% of the variation in LOEUF, and the contribution of CpG density exceeds the individual contributions of exonic and promoter conservation. We leverage this to train a simple and easily interpretable predictive model that out-performs other existing predictors and allows us to classify 1,760 genes – which currently lack reliable LOEUF estimates – as highly LoF-intolerant or not. These predictions have the potential to aid in the interpretation of novel patient variants. Moreover, our results reveal that high CpG density is not merely a generic feature of human promoters, but is preferentially encountered at the promoters of the most selectively constrained genes, calling into question the prevailing view that CpG islands are not subject to selection.

## Introduction

A powerful way of gaining insight into a gene’s contribution to organismal homeostasis is by studying the fitness effect exerted by loss-of-function (LoF) variants in that gene. Fully characterizing this effect is challenging, as it requires estimation of both the selection coefficient for individuals with biallelic LoF variants, as well as the dominance coefficient (Falconer, Mackay, 1996; Fuller et al., 2019). However, recent studies based on the joint processing and analysis of large numbers of exome sequences have developed metrics which serve as approximations to genic LoF-intolerance in humans (Petrovski et al., 2013; Lek et al., 2016; Karczewski et al., 2019). These metrics correlate with several properties indicative of LoF-intolerance (such as enrichment for known haploinsufficient genes; Lek et al. (2016) and Karczewski et al. (2019)), and can substantially help in the assignment of pathogenicity to novel variants encountered in patients as recommended by the American College of Medical Genetics and Genomics (Abou Tayoun et al., 2018).

At the core of all these metrics is a comparison of the observed to the expected number of LoF variants. Hence, genes where the latter is small (e.g. due to small coding sequence length or low mutation rate) will not be amenable to this approach until the sample sizes become much larger than they presently are. Currently in gnomAD, the largest such effort with publicly available constraint data based on 125,748 exomes, approximately 28% of genes lack reliable LoF-intolerance estimates (Karczewski et al., 2019). It has been estimated that even with 500,000 indviduals, the discovery of LoF variants will remain far from saturation, with potentially a sizeable fraction of genes still difficult to ascertain (Zou et al., 2016).

The cardinal feature of highly LoF-intolerant genes, genes depleted of even monoallelic LoF variants in healthy individuals, is dosage sensitivity; a gene copy containing one or more LoF variants produces mRNAs that are typically degraded via nonsense-mediated decay (Lykke-Andersen, Jensen, 2015; Lindeboom et al., 2019). Therefore, the deleterious effects of LoF variants in these genes are often mediated through a reduction of the normal amount of mRNA used for protein production. This in turn, implies that studying the characteristics of regulatory elements controlling the expression of highly LoF-intolerant genes has the potential to yield two important benefits (Han et al., 2018; Wang, Goldstein, 2020). First, it can highlight the features of the most functionally important regulatory elements in the human genome. Second, such features can then provide the basis for predictive models of LoF-intolerance, which can be applied to unascertained genes.

In promoters, one sequence feature that has been extensively studied is CpG density. A large number of mammalian promoters harbor CpG islands (AP Bird, 1987; Deaton, A Bird, 2011), which typically remain constitutively unmethylated in all cell types (Meissner et al., 2008; Straussman et al., 2009). Recently, it has been shown that clusters of unmethylated CpG dinucleotides are recognized by CxxC-domain containing proteins (Lee et al., 2001; Long et al., 2013), thereby facilitating the deposition of transcription-associated marks such as H3K4me3 (Thomson et al., 2010; Clouaire et al., 2012; Wachter et al., 2014). Additionally, there is now evidence that unmethylated CpGs surrounding transcription factor (TF) motifs may contribute to promoter activity by also increasing the probability that the cognate TFs will bind (White et al., 2013; Hartl et al., 2019).

## Results

### Promoter CpG density is strongly and quantitatively associated with downstream gene LoF-intolerance

We discovered a strong relationship between the observed-to-expected CpG ratio (hereafter referred to as CpG density) of a promoter, and LoF-intolerance of the downstream gene (Figure 1a, b); high CpG density is associated with high LoF-intolerance. To establish this, we used the LOEUF metric provided by gnomAD, an updated and more accurate measure of genic LOF-intolerance compared to pLI (Karczewski et al., 2019). LOEUF places human genes on a 0-to-2 continuous scale, with lower values indicating higher LoF-intolerance. Following previous work (Cummings et al., 2019), we classified genes with LOEUF < 0.35 as highly LoF-intolerant. In Karczewski et al. (2019), genes with 10 ≤ expected LoF variants were found to have unreliable LOEUF estimates. Based on additional assessment (Supplemental Figure S1; Methods), we here adopted a more stringent threshold and considered 8,506 genes with ≥ 20 expected LoF variants. We further filtered this set down to 4,743 genes for which we could reliably determine the canonical promoter (Supplemental Figure S2; Methods; Supplemental Figure S3 contains a schematic of our approach to partitioning genes according to the reliability of their LOEUF estimate and promoter annotation).

**Figure 1.**
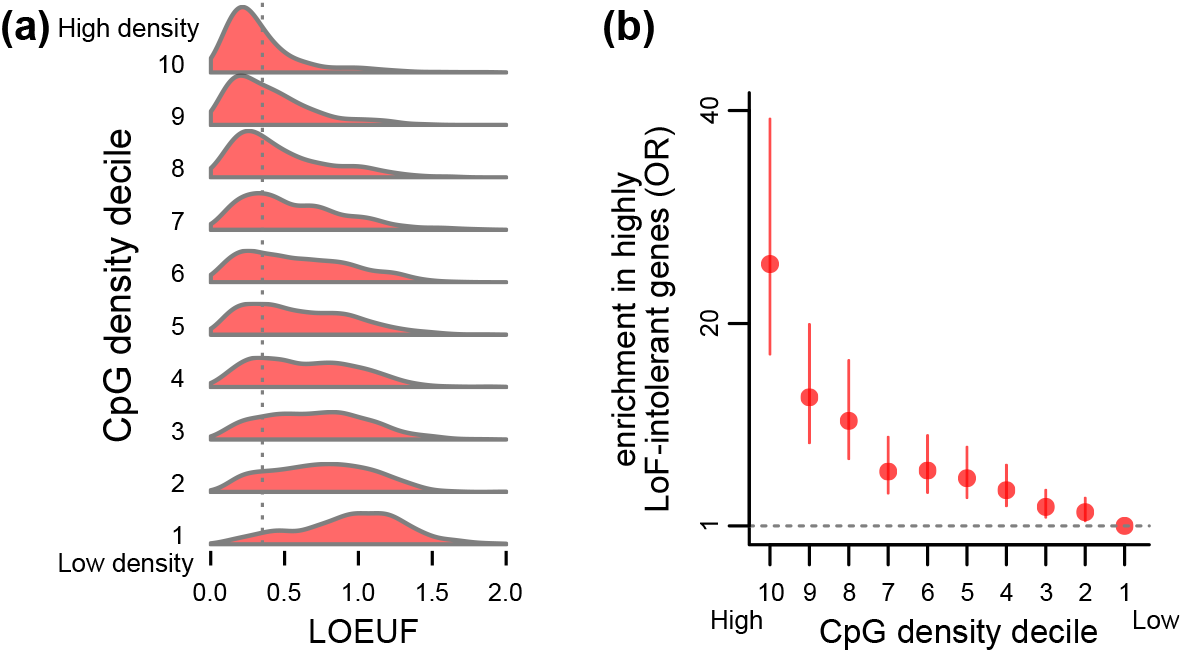
The relationship between promoter CpG density and downstream gene loss-of-function intolerance. **(a)** The distribution of genic LOEUF (as provided by gnomAD) in each decile of promoter CpG density. The vertical line corresponds to the cutoff for highly LoF-intolerant genes (LOEUF < 0.35). **(b)** Odds ratios and the corresponding 95% confidence intervals, quantifying the enrichment for highly LoF-intolerant genes (LOEUF < 0.35) that is exhibited by the set of genes in each decile of promoter CpG density. For each of the other deciles, the enrichment is computed against the 10th decile. The horizontal line corresponds to zero enrichment. In both **(a)** and **(b)**, CpG density deciles are labeled from 1-10 with 1 being the most CpG-poor and 10 the most CpG-rich decile.

When ranked according to the CpG density of their promoter, genes in the top 10% have a 67.2% probability of being highly LoF-intolerant. This in contrast to 7.4% for genes in the bottom 10%, yielding a 25.6-fold enrichment (*p* < 2.2 · 10^−16^; Figure 1b). We note that there is a continuous gradient of enrichment across CpG density deciles (Figure 1b). When splitting genes into just two groups, consisting of those with CpG island-overlapping promoters, and those without, we found that the enrichment for highly LoF-intolerant genes in the CpG-island-overlapping group is markedly weaker (odds ratio = 3.71, *p* < 2.2 · 10^−16^), showing that this dichotomy masks the more continuous nature of CpG density. Finally, regression modeling revealed that CpG density alone can explain 19.3% of the variation in LOEUF (*p* < 2.2 · 10^−16^, *β* = −1.02; (Supplemental Figure S4; Methods), and that its effect on LOEUF is unchanged when accounting for coding sequence length (*p* < 2.2 · 10^−16^, *β* = −1.00).

We emphasize that our result remains pronounced even when we omit the filtering for high-confidence promoters, and merely consider all canonical promoters with ≥ 20 expected LoF variants (*p* < 2.2 · 10^−16^; Supplemental Figure S5). However, the association becomes weaker (14.6-fold enrichment of highly LoF-intolerant genes in the top CpG density decile), underscoring the importance of accurate promoter annotation. We also verified that the exact definition of the promoter (in terms of the size of the interval around the TSS) has only a small impact on strength of the relationship between CpG density and LOEUF (Supplemental Figure S6).

### The association between CpG density and LoF-intolerance is not mediated through expression level or tissue specificity

The more LoF-intolerant a gene is, the more broadly it tends to be expressed across tissues, and at higher levels (Lek et al., 2016; Karczewski et al., 2019). Even though it is well established that promoter CpG density is associated with these two properties as well (Saxonov et al., 2006; Agarwal, Shendure, 2018; Hartl et al., 2019), we found that neither variable explains our result (Figure 2, Supplemental Figure S7). First, after stratifying genes according to either expression level or tissue specificity (using RNA-seq data from the GTEx consortium; Methods), we saw a clear relationship between promoter CpG density and LOEUF within each stratum (Figure 2a, b). Second, the effect of CpG density on LOEUF is almost equally strong when adjusting for either expression level or tissue specificity (regression *β* = −1.00 and −0.85, respectively, *p* < 2.2 · 10^−16^ for both regression models). Third, even the combination of the two expression properties explains less LOEUF variance than CpG density by itself (Figure 2c).

**Figure 2.**
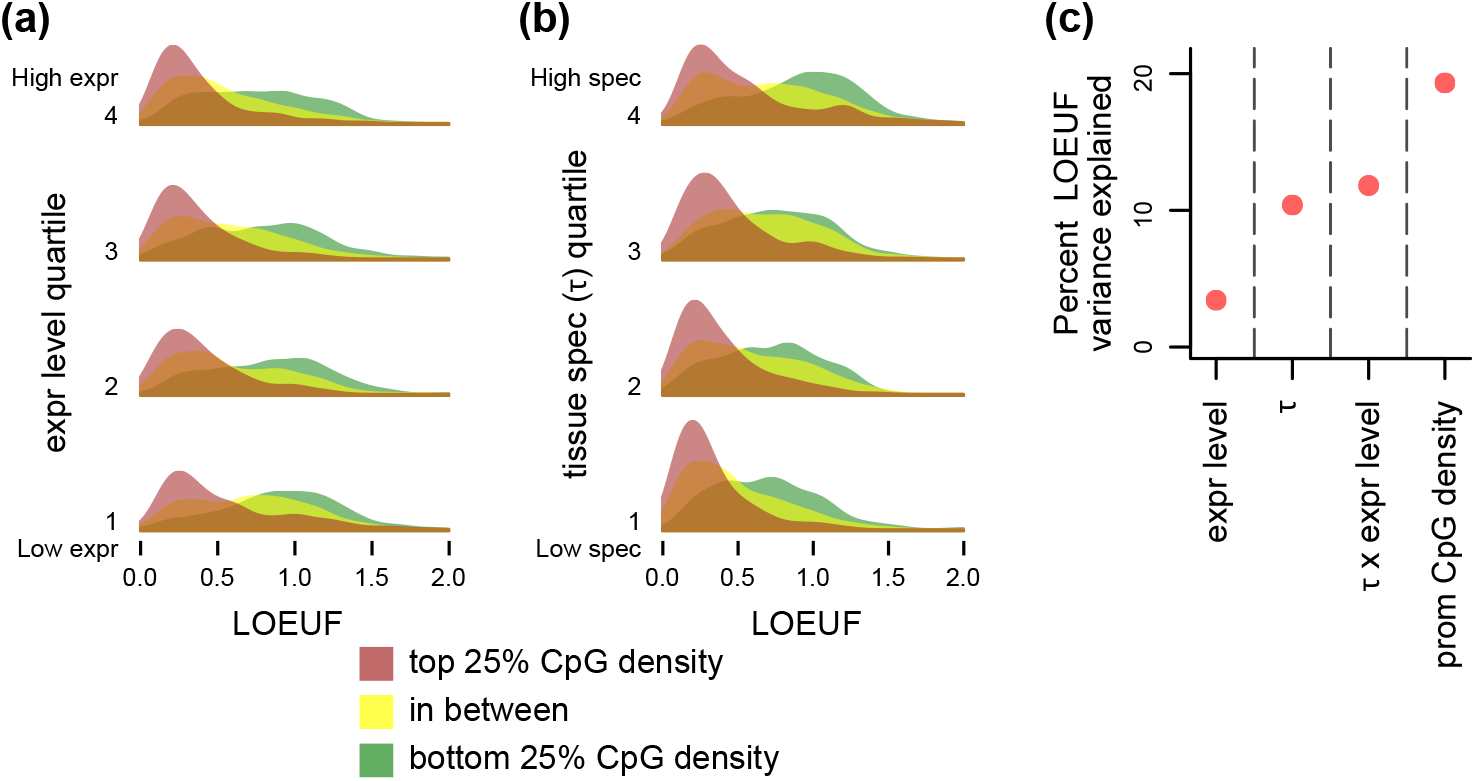
The relationship between promoter CpG density and loss-of-function intolerance conditional on downstream gene expression level and tissue specificity (*τ*). **(a)** The distribution of LOEUF, stratified by promoter CpG density, in each quartile of downstream gene expression level, computed using the GTEx dataset (Methods). **(b)** The distribution of LOEUF, stratified by promoter CpG density, in each quartile of downstream tissue specificity. For each gene, tissue specificity is quantified by *τ*, and is computed using the GTEX dataset (Methods). For both **(a)** and **(b)** quartiles are labeled from 1-4, with 1 being the quartile with the lowest and 4 the quartile with the highest expression/tissue specificity, respectively. **(c)** The percentage of LOEUF variance (adjusted *r*^2^) that is explained by downstream gene expression level, *τ*, the interaction between the two, and promoter CpG density.

### Regulatory factor binding at promoters provides information about LoF-intolerance which adds to CpG density

We next turned our attention to the fraction of LOEUF variation (80.7%) that remains unexplained by CpG density. We hypothesized that part of it might be explained by preferential binding of specific regulatory factors at LoF-intolerant gene promoters. Since a comprehensive assessment of this is currently out of reach (due to the lack of extensive genome-wide binding data for most regulatory factors), we focused on EZH2 as a proof-of-principle. EZH2 is a relatively well-characterized histone methylstransferase that specifically localizes to CpG islands of non-transcribed genes (Figure 3a, Supplemental Figure S8; Riising et al. (2014) and Berrozpe et al. (2017)).

**Figure 3.**
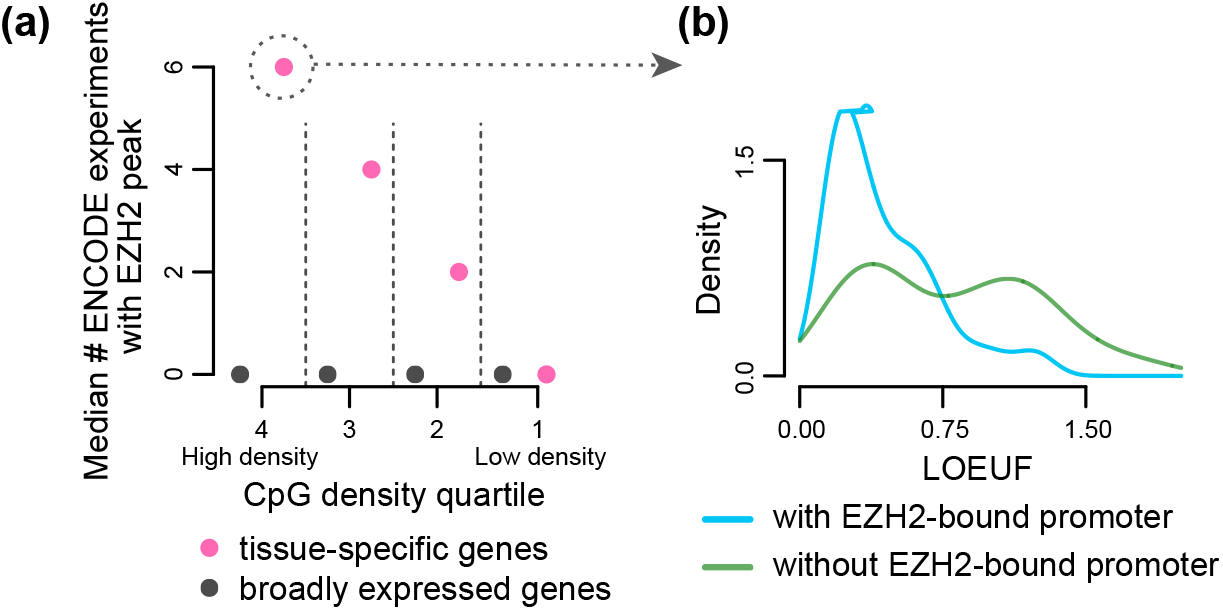
The loss-of-function intolerance of tissue-specific genes conditional on high promoter CpG-density and promoter EZH2 binding. **(a)** The median number of ENCODE ChIP-seq experiments (out of 14 total) where an EZH2 peak is detected, shown separately for tissue-specific (*τ* > 0.6) and broadly expressed (*τ* < 0.6) genes, within each quartile of promoter CpG density. The quartiles are labeled from 1-4, with 1 being the most CpG-poor and 4 the most CpG-rich. **(b)** The LOEUF distributions of tissue-specific genes with high-CpG-density (top 25%) promoters, stratified according to whether their promoters show EZH2 peaks in at least 2 ENCODE experiments, or in less than 2 experiments.

We discovered that tissue-specific genes with CpG-dense and EZH2-bound promoters (EZH2 binding in at least 2 ENCODE experiments) have lower LOEUF compared to their EZH2-unbound counterparts (Figure 3b; regression *β* = −5.66, *p* = 5.21 · 10^−8^, for the interaction between CpG density and EZH2 binding, conditional on tissue specificity *τ* > 0.6). In this subset of promoters, the interaction of EZH2 binding with CpG density explains an additional 27.1% of LOEUF variance on top of what CpG density explains (2.1%).

### Promoter CpG density with promoter and exonic across-species conservation can collectively predict LoF-intolerance with high accuracy

We then sought to develop a predictive model for LoF-intolerance, with the goal of providing high-confidence predictions for the 2,430 genes with currently unreliable LOEUF scores and reliable promoter annotation. Specifically, we aimed to classify genes as highly LoF-intolerant (LOEUF< 0.35) or not.

To build our model, we first separately computed the promoter and exonic across-species conservation for each gene (using the PhastCons score; Methods), and asked if they provide information about LOEUF complementary to CpG density. We found this to be true (Figure S9c and Supplementary Figure S9a,b); notably, CpG density explains at least as much LOEUF variance as exonic or promoter conservation (Figure 4a). When all three metrics are combined, 33.4% of the total LOEUF variation is explained (Figure 4a). We note that while EZH2 binding explains a substantial amount of LOEUF variance when considering tissue-specific genes with high-CpG-density promoters, these are a small subset. Hence, inclusion of this feature only minimally increases the overall explained variance (0.4% increase). We therefore settled on training a logistic regression model with CpG density, and promoter/exonic conservation as three linear predictors. As our training set we used 3,000 genes, randomly selected from the 4,743 with high-confidence LOEUF estimates.

**Figure 4.**
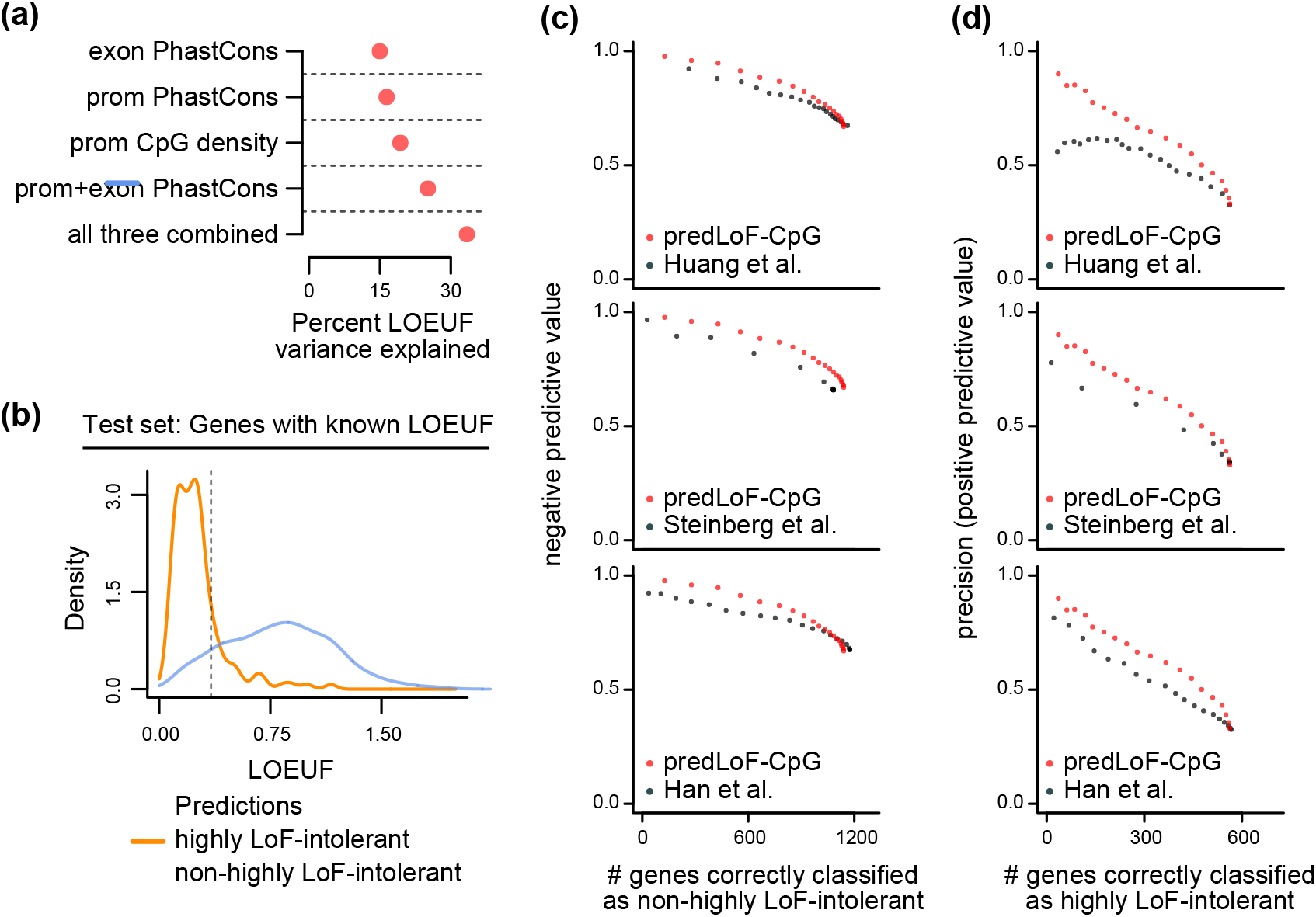
Training and assessing predLoF-CpG: a predictor of loss-of-function intolerance based on CpG density. **(a)** The percentage of LOEUF variance (adjusted *r*^2^) that is explained by CpG density, exonic or promoter conservation, and their combinations. **(b)** The out-of-sample performance of predLoF-CpG. Shown are the LOEUF distributions of 1,743 genes belonging to the holdout test set (which consists of genes with reliable LOEUF estimates), stratified according to their classification as highly LoF-intolerant or not. The dashed vertical line corresponds to the cutoff for highly LoF-intolerant genes (LOEUF < 0.35). **(c)** The precision (y axis, left column) and negative predictive value (y axis, right column) plotted against the number of correctly classified genes (x axis), for different predictors of loss-of-function intolerance. Each point corresponds to a threshold. The thresholds span the [0,1] interval, with a step size of 0.05. We note that because we are using two classification thresholds, a ROC curve would not be an appropriate evaluation metric here.

Our predictor, which we called predLoF-CpG (predictor of LoF-intolerance based on CpG density) showed strong out-of-sample performance on the test set of the remaining 1,743 genes. The precision (positive predictive value) was 82.6% at the 0.75 prediction probability cutoff, and the negative predictive value was 88.4% at the 0.25 cutoff (Figure 4b); 144 genes were predicted to be highly LoF-intolerant, 753 were predicted as non-highly LoF-intolerant, and 806 (47.3%) were left unclassified. We chose to use two thresholds instead of one, at the expense of leaving a fraction of genes unclassified, since this endows our predictor with precision and negative predictive value high enough to be useful in the clinical setting. We note that our predictive accuracy is comparable to that of widely adopted tools for predicting damaging missense variants (Sim et al., 2012; Adzhubei et al., 2013). Further examining our out-of-sample classifications, we found that a) the genes falsely predicted as highly LoF-intolerant had a median LOEUF of 0.49, indicating that at least half of them are very LoF-intolerant despite not exceeding the 0.35 cutoff, and b) the genes correctly predicted as non-highly LoF intolerant had a median LOEUF of 0.86, suggesting that at least half of them are likely to tolerate biallelic inactivation as well (Figure 4b).

Regardless of the choices for the two classification thresholds, predLoF-CpG outperforms all of the previously published predictors of LoF-intolerance (Figure 4c). Specifically, all models have comparable and high negative predictive value, with ours being slightly superior (Figure 4c). However, within a range of thresholds that yield high precision, as would be required for use in clinical decision making, predLoF-CpG provides clear gain versus the rest (Figure 4d, upper left area of left column plots). As an additional evaluation, we found that predLoF-CpG is capable of explaining a greater proportion of out-of-sample LOEUF variance compared to the other three (Supplemental Figure S10).

Finally, we mention GeVIR, a recently developed metric (primarily for intolerance to missense, but also useful for LoF variation; Abramovs et al. (2020)) which identifies regions depleted of protein-altering variation (Havrilla et al., 2019), and weights these regions by conservation within each gene. As expected given its dependency on observed variation, GeVIR exhibits substantial correlation with the expected number of LoF variants (Spearman correlation = 0.42 vs 0.26 for predLoF-CpG). This limits its applicability to genes with unreliable LOEUF, even though the weighting step slightly alleviates this issue compared to LOEUF (Spearman correlation = 0.49).

### 32.5% of currently unascertained genes in gnomAD receive high-confidence predictions by predLoF-CpG

We applied predLoF-CpG to genes with unreliable LOEUF estimates in gnomAD. After filtering for these with high-confidence promoter annotation, we retained 2,430 (out of 5,413). Of these, 104 were classified as highly LoF intolerant, 1,656 as non-highly LoF intolerant and 670 were left unclassified (Supplemental Table 1). We first examined the ratio of observed-to-expected LoF variants in these genes. Even though these point estimates are uncertain, there is a clear difference in the distribution of the point estimates between genes we classify as highly LOF intolerant (median = 0.14) and those as not (median = 0.70), with the difference being in the expected direction (Figure 5a; Wilcoxon test, *p* < 2.2 · 10^−16^).

**Figure 5.**
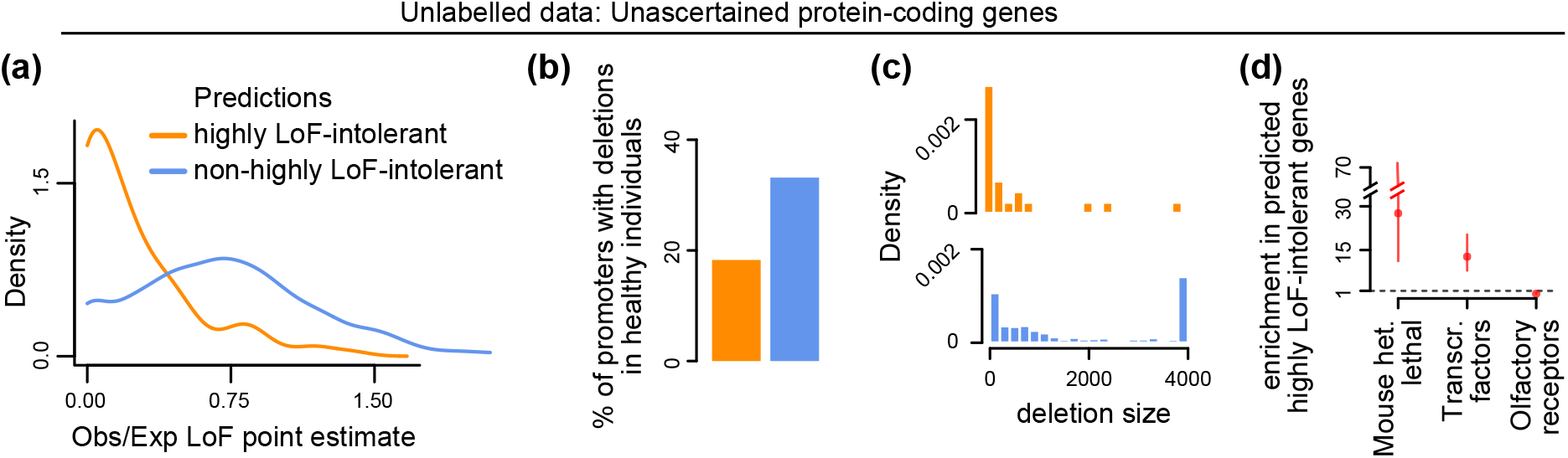
Using predLoF-CpG to classify currently unascertained genes as highly loss-of-function intolerant or not. **(a)** The distribution of point estimates of the observed/expected proportions of LoF variants. Genes are stratified according to their classification as highly LoF-intolerant or not. **(b)** The proportion of promoters which harbor deletions in a sample of 14,891 healthy individuals. Promoters are stratified according to downstream gene classification as highly LoF-intolerant or not. **(c)** The distribution of the size of deletions harbored by promoters in a sample of 14,891 healthy individuals. Promoters are stratified according to downstream gene classification as highly LoF-intolerant or not. **(d)** Odds ratios and the corresponding 95% confidence intervals quantifying the enrichment for genes in each of the x-axis groups that is exhibited by genes predicted as highly LoF-intolerant by predLoF-CpG. The enrichment is computed against genes predicted as non-highly LoF-intolerant. The horizontal line at 1 corresponds to zero enrichment.

Next, to provide orthogonal support for our predictions, we leveraged a set of 175,716 deletions detected in 14,891 healthy individuals using whole-genome sequencing (Methods) (Collins et al., 2019). We reasoned that LoF-intolerant gene promoters should be depleted of such deletions; when they do harbor deletions, these should be small. By only considering promoters, we ensured that our assessment is not dependent on gene length, which confounds LOEUF estimation. Using the 4,743 genes with high-confidence LOEUF (from the training and test sets), we first observed that low LOEUF is indeed associated with the presence of both fewer (*p* = 2.39 · 10^−15^) and smaller (*p* < 2.2 · 10^−16^) promoter deletions (Supplemental Figure S11a, b), showing that this is a legitimate assessment strategy. Turning to our predictions, we found the same: genes predicted to be highly LoF-intolerant are less likely to contain deletions in their promoters compared to genes classified as non-highly LoF-intolerant (Figure 5b; probability of overlapping at least one deletion = 0.18 vs 0.33, permutation one-sided *p* = 4 · 10^−4^ after 10,000 permutations); when such deletions are observed, they tend to be much smaller (Figure 4c; median size = 129 vs 1092; Wilcoxon test, *p* = 4.49 · 10^−5^).

Finally, we found that our predictions are in strong agreement with what would be expected based on known mouse phenotypes, and membership in specific gene classes (Figure 5d). First, the predicted highly LoF-intolerant genes show a 27.6-fold enrichment for genes heterozygous lethal in mouse (*p* = 1.03 · 10^−12^), when compared against those predicted as non-highly LoF-intolerant. Second, they exhibit a 12.7-fold enrichment for transcription factors (*p* < 2.2 · 10^−16^), consistent with the known dosage sensitivity of these genes (Jimenez-Sanchez et al., 2001; JG Seidman, C Seidman, 2002; Boukas et al., 2019). Third, they show a total depletion (odds ratio = 0) of olfactory receptor genes (*p* = 2.5 · 10^−5^).

### predLoF-CpG reclassifies 101 genes with expected LoF variants between 10 and 20 as highly LoF-intolerant

In our analyses so far, we have ignored 3,440 genes with expected LoF variants between 10 and 20. Even though in Karczewski et al. (2019) these were treated as having reliable LOEUF estimates, our assessment suggests that lack of power can affect whether they are categorized as highly LoF intolerant or not (Supplementary Figure S1, Methods). After filtering for reliable promoter annotation, we applied predLoF-CpG to 2,772 genes, and obtained high-confidence classifications for 1,675. For the great majority (93.9%), we agree with the classification obtained by purely considering whether their LOUEF is < 0.35. However, we observed 101 genes that were classified as highly LoF-intolerant by predLoF-CpG but had LOEUF ≥ 0.35, a number not explained by the false positive rate of our predictor (Supplemental Table 2). 75% of these genes have an observed/expected LoF point estimate of 0.31, suggesting that they are indeed highly LoF-intolerant, but do not exceed the required LOEUF threshold because of inadequate power. Therefore, when interpreting LoF variants in these genes, we suggest that both LOEUF as well as predLoF-CpG are taken into account.

## Discussion

Our study reveals that: a) there exists a strong, widespread coupling between promoter CpG density and downstream gene LoF-intolerance in the human genome, and b) this coupling can be exploited to predict LoF-intolerance for almost 2000 genes that are otherwise largely intractable with current sample sizes. Our predictions for these genes (which we make available in Supplemental Table 1) can inform research into novel disease candidates and now become incorporated in the clinical genetics laboratory setting. Similarly to existing tools for missense variants (Sim et al., 2012; Adzhubei et al., 2013), they can provide corroborating evidence during the evaluation of the pathogenicity of LoF variants harbored by patients, as recommended by the American College of Medical Genetics and Genomics (Abou Tayoun et al., 2018).

In terms of understanding the regulatory architecture of the genome, our findings extend decades of work (AP Bird, 1987; Deaton, A Bird, 2011) to show that high CpG density is not just a prevalent feature of many promoters, but is preferentially marking the promoters of the most selectively constrained genes. We believe this casts doubt on the prevailing view that CpG islands are not under selection (Cohen et al., 2011), although we note that our current results are correlative in nature.

If promoter CpG density is indeed under selection, its presence at LoF-intolerant gene promoters has to be advantageous, which raises the question of the underlying biological mechanism. Our findings suggest that this mechanism is not related to the high and constitutive expression that LoF-intolerant genes typically exhibit. An intriguing possibility has been recently raised by single-cell expression measurements showing that promoter CpG islands are associated with reduced expression variability (Morgan, Marioni, 2018). We hypothesize that this decreased variability is beneficial for many processes where LoF-intolerant genes are known to play central roles, such as neurodevelopment (Fahrner, Bjornsson, 2019).

Our work represents an attempt at deciphering the link between regulatory element characteristics, and the LoF-intolerance of the genes they control. The fact that taking promoter EZH2 binding into account improves our ability to recognize LoF-intolerant genes on top of CpG density, implies that this mapping can be learned with even greater accuracy by incorporating information about other regulatory factors as well. However, a current barrier to achieving this is the relative paucity of genome-wide binding data across the full repertoire of transcription factors: the human genome encodes approximately 1500 transcription factors (Vaquerizas et al., 2009; Barrera et al., 2016; Lambert et al., 2018) and at least 300 epigenetic regulators (Boukas et al., 2019). In contrast to these numbers, currently ENCODE has profiled only 330 regulatory factors in K562 cells, the most extensively characterized cell line.

It is also natural to consider moving beyond promoters to other regulatory elements. An initial step in this direction has recently been taken in Wang, Goldstein (2020), motivated by work in *Drosophila* showing that developmentally important genes can have multiple redundant enhancers (Perry et al., 2010; Frankel et al., 2010). While this “enhancer domain score” was not designed to capture LoF-intolerance and has poor association with LOEUF (adjusted *r*^2^ = 0.03), it has been shown to have some predictive capacity for human disease genes, especially those with a developmental basis.

In summary, our study shows the existence of a strong and widespread association between promoter CpG density and genic LoF-intolerance, and leverages this relationship to predict LoF-intolerance for unascertained genes.

## Methods

### Selecting transcripts with high-confidence loss-of-function intolerance estimates

In total, gnomAD (Karczewski et al., 2019) provides LoF-intolerance estimates for 79,141 human proteincoding transcripts (hereafter referred to as trancripts) labeled with ENSEMBL identifiers, of which 19,172 are annotated as canonical. For each transcript, these LoF-intolerance estimates consist of the point estimate of the observed/expected number of LoF variants, as well as a 90% confidence interval around it. The upper bound of this confidence interval (LOEUF) is the suggested metric of LoF-intolerance (Karczewski et al., 2019). For any given transcript, the ability to reliably estimate LOEUF is directly related to the expected number of LoF variants; when that expected number is small, there is uncertainty around the point estimate (and thus a large LOEUF value), because it is not possible to determine whether an observed depletion of LoF variants is due to negative selection against these variants in the population, or due to inadequate sample size. Therefore, for transcripts with high-confidence LOEUF values, there should be a strong positive correlation between the point estimate and LOEUF; in constrast, low-confidence LOEUF transcripts will have small point estimates coupled with large LOUEF values.

Based on this assessment, and consistent with Karczewski et al. (2019), we considered transcripts with ≤ 10 expected LoF variants to have unreliable LOEUF (34,232 out of 79,141 total transcripts; 5,413 out of 19,172 canonical transcripts; Supplemental Figure S1). Throughout the text, we refer to the genes encoding for these transcripts as “unascertained”.

Even though in Karczewski et al. (2019) most of the analyses were performed using transcripts with > 10 expected LoF variants, we saw that, with increasing expected number of LoF variants, there was a non-negligible increase in the probability (conditional on a given point estimate) of a transcript belonging in the highly LoF-intolerant category (LOEUF < 0.35). We thus adopted a more stringent criterion, and considered transcripts with ≥ 20 expected LoF variants (25,474 out of 79,141 total transcripts; 8,506 out of 19,172 canonical transcripts; Supplemental Figure S1) to have high-confidence LOEUF. After further filtering based on promoter annotation (see the section “Annotating canonical promoters in the human genome”), these are the transcripts we used to establish the association between promoter CpG density and LOEUF, and to train predLoF-CpG.

### Selecting transcripts with high-confidence annotations in GENCODE v19

gnomAD supplies LOEUF estimates for 79,141 transcripts in GENCODE v19. However, we conducted our analyses at the gene level, based on the following reasoning: typically, transcripts from the same gene have overlap in their coding sequence, which makes it hard to disentangle their LOEUF estimates. For example, a transcript whose loss does not have severe phenotypic consequences, and therefore its promoter does not contain informative features, may still have low LOEUF merely because it overlaps with a different transcript of the same gene.

For each gene, GENCODE labels a single transcript as canonical, and recognizes the difficulty of accurately annotating transcriptional start sites (TSS’s) (Harrow et al., 2012). We manually inspected GENCODE’s choices of canonical transcripts, and found some problematic cases. An illustrative example is *KMT2D* (Supplemental Figure S12). First, even though this gene is broadly expressed across tissues in GTEx, its canonical promoter shows POLR2A (the major subunit of RNA PolII complex) ChIP-seq peaks in only 4 ENCODE experiments (out of 74 total). Even though there does exist a non-canonical transcript whose promoter has POLR2A signal in 59 experiments, as would expected for a broadly expressed gene, that non-canonical transcript has an unusually short coding sequence, which does not even encode for the catalytic SET domain. In this particular case, we reasoned that the 5’ UTR of the canonical transcript needs to be extended up until the TSS of the non-canonical transcript. Such an annotation would also be consistent with the annotation of the mouse ortholog. Importantly, if this annotation error is ignored, it is impossible to select a *KMT2D* transcript with accurate estimates of both LOEUF and promoter CpG density.

With this example in mind, we developed an empirical approach to only retain transcripts with high-confidence GENCODE annotations in our analysis. First, we defined promoters as 4kb elements centered around the transcriptional start site (TSS). We then leveraged the main hallmark of transcriptional initiation at protein-coding gene promoters: binding of the RNA PolII complex, the major subunit of which is POLR2A (Wintzerith et al., 1992; Mita et al., 1995). We used data from ENCODE (ENCODE Project Consortium, 2012) on the genome-wide binding locations of POLR2A from 74 ChIP-seq experiments on several cell lines, originating from diverse human tissues (see “POLR2A ENCODE ChIP-seq data” section below).

As expected, we observed that genes that are broadly expressed across the 53 different tissues in GTEx (*τ* < 0.6; see “GTEx expression data” section below) tend to have promoters with POLR2A ChIP-seq peaks in multiple experiments, while the opposite is true for genes expressed in a restricted number of tissues (*τ* > 0.6, Supplemental Figure S2c). However, as in the *KMT2D* example above, we also observed genes with broad expression and very low binding of POLR2A at their canonical promoter (Supplemental Figure S2c), and a few genes with restricted expression but POLR2A peaks at their canonical promoter in multiple experiments (Supplemental Figure S2c), raising our suspicion that these reflect inaccurate annotation of the canonical TSS.

Therefore, we required that the canonical promoter of a broadly expressed gene exhibits POLR2A peaks in multiple ENCODE experiments, and the canonical promoter of a gene with restricted expression exhibits POLR2A peaks only in a small number of ENCODE experiments. As additional layers of evidence for canonical promoters, we used the presence of CpG islands, which are known markers of promoters in mammalian genomes (AP Bird, 1987; Deaton, A Bird, 2011), as well as the concordance between the human TSS coordinate and the TSS coordinate of a mouse ortholog transcript (when the latter is mapped onto the human genome).

Specifically, we first excluded genes on the sex chromosomes, since, due to X-inactivation in females and hemizygosity in males, LoF-intolerance estimates have different interpretion in these cases. This gave us 17,657 genes with at least one canonical transcript, of which 17,359 had expression measurements in GTEx. We then applied the following criteria (when none of the criteria were satisfied, we entirely discarded the gene):

#### Criterion 1

The gene is broadly expressed (*τ* < 0.6) and the canonical promoter has a POLR2A peak in more than 35 ENCODE experiments.

We found 7,250 cases satisfying this criterion, and kept the canonical promoter annotation.

#### Criterion 2

The gene is broadly expressed (*τ* < 0.6), the canonical promoter has a POLR2A peak in less than 10 ENCODE experiments, and there is an alternative promoter with POLR2A peaks in more than 35 experiments.

We found 218 cases satisfying this criterion (Supplemental Figure S2d), and classified the alternative promoter as the canonical (all such cases are provided in Supplemental Table 3). When there were more than one alternative promoters satisfying our requirement, we distinguished the following subcases:

a. If none of these alternative promoters overlapped a CpG island, we classified the promoter corresponding to the transcript with the greater number of expected LoF variants as the canonical.
b. If exactly one of these alternative promoters overlapped a CpG island, we classified that promoter as the canonical.
c. If more than one of these alternative promoters overlapped a CpG island, we classified the promoter that, among the CpG-island-overlapping promoters, had the greatest number of expected LoF variants as the canonical.

For our subsequent analyses, we used the LOEUF value of the newly annotated canonical promoter.

#### Criterion 3

The gene is not broadly expressed (*τ* > 0.6), the canonical promoter has a POLR2A peak in less than 10 ENCODE experiments, and overlaps a CpG island.

We found 1,862 cases satisfying this criterion, and kept the canonical promoter annotation.

#### Criterion 4

The gene is not broadly expressed (*τ* > 0.6), the canonical promoter has a POLR2A peak in less than 10 ENCODE experiments, none of the promoters corresponding to the gene overlap a CpG island, and there is a mouse ortholog TSS in RefSeq no more than 500bp away from the canonical human TSS.

We found 3,049 cases satisfying this criterion, and kept the canonical promoter annotation.

#### Criterion 5

The gene is not broadly expressed (*τ* > 0.6), the canonical promoter has a POLR2A peak in less than 10 ENCODE experiments, none of the promoters corresponding to the gene overlap a CpG island, there is no mouse ortholog TSS in RefSeq, and there are no alternative transcripts with different TSS coordinates.

We found 1,411 cases satisfying this criterion, and kept the canonical promoter annotation.

The promoters selected from the above 5 criteria along with their coordinates are provided in Supplemental Table 4.

Finally, regarding coding sequence annotations, errors such as the one in *KMT2D* described at the beginning of the section are difficult to systematically detect and correct, and our manual inspection suggested that they are also less frequent. We chose to entirely discard cases where:

a. the trascript we had selected after promoter filtering had ≤ 10 expected LoF variants (placing the gene into the “unascertained” category), and
b. there was an alternative transcript that had longer coding sequence and ≥ 20 more expected LoF variants compared to the one our procedure selected.

This approach removes *KMT2D* and 14 more potentially problematic cases such as *ZNF609*.

### Calculating the CpG density of a promoter

Using the BSgenome.Hsapiens.UCSC.hg19 R package, we obtained the sequence of each promoter with the get-Seq function. We then calculated its CpG density using the definition of the observed-to-expected CpG ratio in Gardiner-Garden, Frommer (1987), applied to the entire 4 kb sequence (that is, without using sliding windows).

### The impact of promoter definition

There is currently no single accepted definition of a promoter in terms of the size of the interval around the TSS. We therefore examined how this parameter affects the relationship between CpG density and LOEUF, and found its impact to be small for 5 sensible choices (Supplemental Figure S6a,b).

### Overlapping promoters

When defining the set of genes with high-confidence LOEUF estimates, we excluded genes whose promoters overlapped promoters of genes with less than 20 expected LoF variants, but whose observed/expected LoF point estimate was suggestive of LoF-intolerance (< 0.5). In cases of overlapping promoters with both genes having ≥ 20 expected LoF variants, we kept the promoter corresponding to the gene with the lowest LOEUF. In cases of overlapping promoters with both genes having at ≤ 10 expected LoF variants, we kept the promoter with the highest CpG density. Finally, when defining the set of unascertained genes, we excluded genes whose promoters overlapped promoters of genes with greater than 10 expected LoF variants, unless there was strong evidence that these were LoF-tolerant (observed/expected LoF point estimate > 0.8 and at least 20 expected LoF variants.)

We recognize however, that in cases where promoters overlap, the predictions are potentially informative not only for the gene whose promoter was ultimately used, but also for the genes with overlapping promoters. In addition, in cases of genes predicted as highly LoF-intolerant, these predictions might also have been influenced by the overlapping promoter (there are only 3 potential such cases). With that in mind, in Supplemental Table 1, we provide such information under the column “other genes with overlapping promoter”.

### Promoters in subtelomeric regions

It is known that subtelomeric regions are rich in CpG islands, which are however different than those in the rest of the genome, in that they appear in clusters, and their CpG-richness is driven mainly by GC-biased gene conversion (Cohen et al., 2011). We thus excluded promoters residing in subtelomeric regions (defined as 2 Mb on each of the two chromosomal ends of each chromosome) from our analyses.

### ENCODE ChIP-seq data

We used the rtracklayer R package to download the “wgEncodeRegTfbsClusteredV3” table from the “Txn Factor ChIP” track, as provided by the UCSC Table Browser for the hg19 human assembly. We then restricted to peak clusters corresponding to POLR2A. This gave us a set of genomic intervals, each of which has been derived from uniform processing of 74 POLR2A ChIP experiments on 32 distinct cell lines (some cell lines were represented by more than one experiments). Each genomic interval was associated with a single number, which ranged from 0 to 74 and indicated the number of ChIP experiments where a peak was detected at that interval. The EZH2 data were downloaded in an identical manner.

### GTEx expression data

We used the GTEx portal to download a matrix with the gene-level TPM expression values from the v7 release, derived from RNA-seq expression measurements from 714 individuals, spanning 53 tissues. (GTEx Consortium, 2017).

As the metric of tissue specificity for a given gene, we used *τ*, which has been shown to be the most robust such measure when benchmarked against alternatives (Kryuchkova-Mostacci, Robinson-Rechavi, 2017). To calculate *τ*, we first computed the gene’s median expression across individuals, within each tissue. Since it has been shown that the transcriptomic profiles of the different brain regions are very similar, with the exception of the two cerebellar tissues (GTEx Consortium, 2015), which are similar to one-another, we aggregated the median expression of each gene in the different brain regions into two “meta-values”. One meta-value corresponded to the median of its median expression in the two cerebellar tissues, and the other to the median of its median expression in the other brain regions. We then formed a matrix where rows corresponded to genes, and columns to tissues, with one column for the across-brain-regions meta-value and another for the across-cerebellar-tissues meta-value; the entries in the matrix were log2(TPM + 1) median expression values. Finally, for each gene, *τ* was calculated as described in (Kryuchkova-Mostacci, Robinson-Rechavi, 2017).

For our analyses of the association between promoter CpG density and expression level, we used the median (across individuals) expression (log_2_(TPM + 1)), computed for the tissue where the gene had the maximum median expression.

### TSS coordinates of mouse orthologs

We used the biomaRt R package to obtain a list of mouse-human homolog pairs, using the human Ensembl gene IDs as the input. For this query, we set the ‘mmusculus homolog orthology confidence’ parameter equal to 1 (indicating high-confidence homolog pairs). Then, for each of the mouse homolog Ensembl IDs, we retrieved the RefSeq mRNA IDs, again with biomaRt. We discarded cases where the same RefSeq mRNA ID was associated with more than one Ensembl gene IDs. We then used the rtracklayer R package to download the “xenoRefGene” UCSC table, from the “Other RefSeq” track, containing the TSS coordinates for each of the mouse RefSeq trancripts.

### Across-species conservation quantification

For each nucleotide, we quantified conservation across 100 vertebrate species using the PhastCons score (Siepel et al., 2005), obtained with the phast-Cons100way.UCSC.hg19 R package. The PhastCons score ranges from 0 to 1 and represents the probability that a given nucleotide is conserved. As the promoter PhastCons score for a given gene, we computed the average PhastCons of all nucleotides in the 4kb region centered around the TSS. As the exonic PhastCons for a given gene, we pooled all nucleotides belonging to the coding sequence of the gene (that is, excluding the 5’ and 3’ UTRs), and computed their average PhastCons.

### Previously published LoF-intolerance predictions

The updated version of the score of Huang et al. (2010) was downloaded from the DECIPHER database (https://decipher.sanger.ac.uk/files/downloads/HI_Predictions_Version3.bed.gz; accessed November 2019). The scores of Steinberg et al. (2015) and Han et al. (2018) were downloaded from the Supplemental materials of the respective publications. In our comparison we did not include HIPred (Shihab et al., 2017), since it only provides binary haploinsufficiency predictions for a small number of genes.

### Structural variation data

We used the gnomAD browser to download a bed file (https://storage.googleapis.com/gnomad-public/papers/2019-sv/gnomad_v2.1_sv.sites.bed.gz.tbi) containing the coordinates and characteristics of structural variants in gnomAD v2. We then restricted to deletions that passed quality control (“FILTER” column value equal to “PASS”). Subsequently, we excluded deletions that overlapped more than one of our high-confidence promoters, in order to avoid ambiguous links between deletions and genes.

### Gene catalogs

The following gene catalogs were used for Figure 5d:

a. 404 heterozygous lethal genes in mouse from https://github.com/macarthur-lab/gnomad_lof/blob/master/R/ko_gene_lists/list_mouse_het_lethal_genes.tsv (see the supplemental material of Karczewski et al. (2019) for details on obtaining this set). We mapped these genes to their human homolog ensembl ids with the biomaRt R package using the “mgi symbol” filter, keeping only pairs with the ‘mmusculus homolog orthology confidence’ parameter equal to 1. This yielded a total of 390 human homologs.
b. 1,254 high-confidence transcription factor genes from Barrera et al. (2016)
c. 371 olfactory receptor genes from https://github.com/macarthur-lab/gene_lists/blob/master/lists/olfactory_receptors.tsv.

### Enrichment quantification

All enrichment point estimates in the text correspond to odds ratios, and the associated p-values were calculated using Fisher’s exact test (two-sided) with the “fisher.test” function in R.

### Code

Code used in this manuscript is available at https://github.com/hansenlab/lof_prediction_paper_repro.

## Supporting information

SuppTable1

SuppTable2

SuppTable3

SuppTable4

## Funding

Research reported in this publication was supported by the National Institute of General Medical Sciences of the National Institutes of Health under award number R01GM121459. LB was supported by the Maryland Genetics, Epidemiology and Medicine (MD-GEM) training program, funded by the Burroughs-Wellcome Fund. HTB received support from the Louma G. Foundation.

## Conflict of Interest

HTB is a paid consultant for Millennium Pharmaceuticals, Inc.

## Supplementary Materials

### Contents

1. **Supplemental Table 1**: predLoF-CpG predictions for genes unascertained in gnomAD. Prediction probabilities are provided in the “prediction probability of high LoF intolerance by predLoF-CpG” column. Probabilities > 0.75 correspond to genes predicted as highly LoF-intolerant, and probabilities < 0.25 to genes predicted as non-highly LoF-intolerant. ENSEMBL gene/transcript ids and coordinates of the promoters used for prediction are also provided; all coordinates refer to hg19.
2. **Supplemental Table 2**: Similar to Supplemental Table 1, but for 101 genes with expected LoF variants between 10 and 20 that were classified as highly LoF-intolerant by predLoF-CpG but had LOEUF ≥ 0.35.
3. **Supplemental Table 3**: Promoter coordinates for cases where our promoter filtering procedure selected a non-canonical promoter. The table contains the promoter coordinates and transcript ENSEMBL ids of both the canonical, as well as the alternative transcript that was selected. All coordinates refer to hg19.
4. **Supplemental Table 4**: Promoter coordinates for 11,059 transcripts where our filtering procedure selected a reliable promoter.
5. **Supplemental Figures S1-S12**.

### Supplemental Figures

**Supplemental Figure S1.**
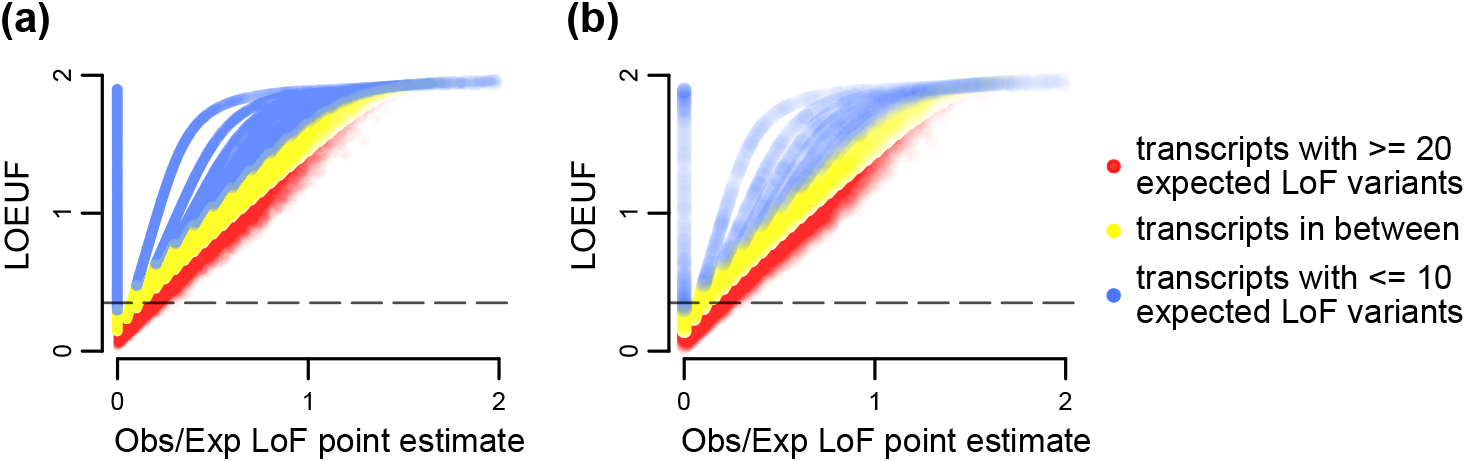
Assessing the reliability of LOEUF estimates. Scatterplots of the point estimates of the observed/expected proportion of loss-of-function variants (x axis), against LOEUF (y axis; defined as the upper bound of the 90% confidence interval around the point estimate). Each point corresponds to a transcript. The horizontal line corresponds to the 0.35 cutoff for highly LoF-intolerant genes. Shown for: **(a)** all transcripts, and **(b)** canonical transcripts only (based on GENCODE annotation).

**Supplemental Figure S2.**
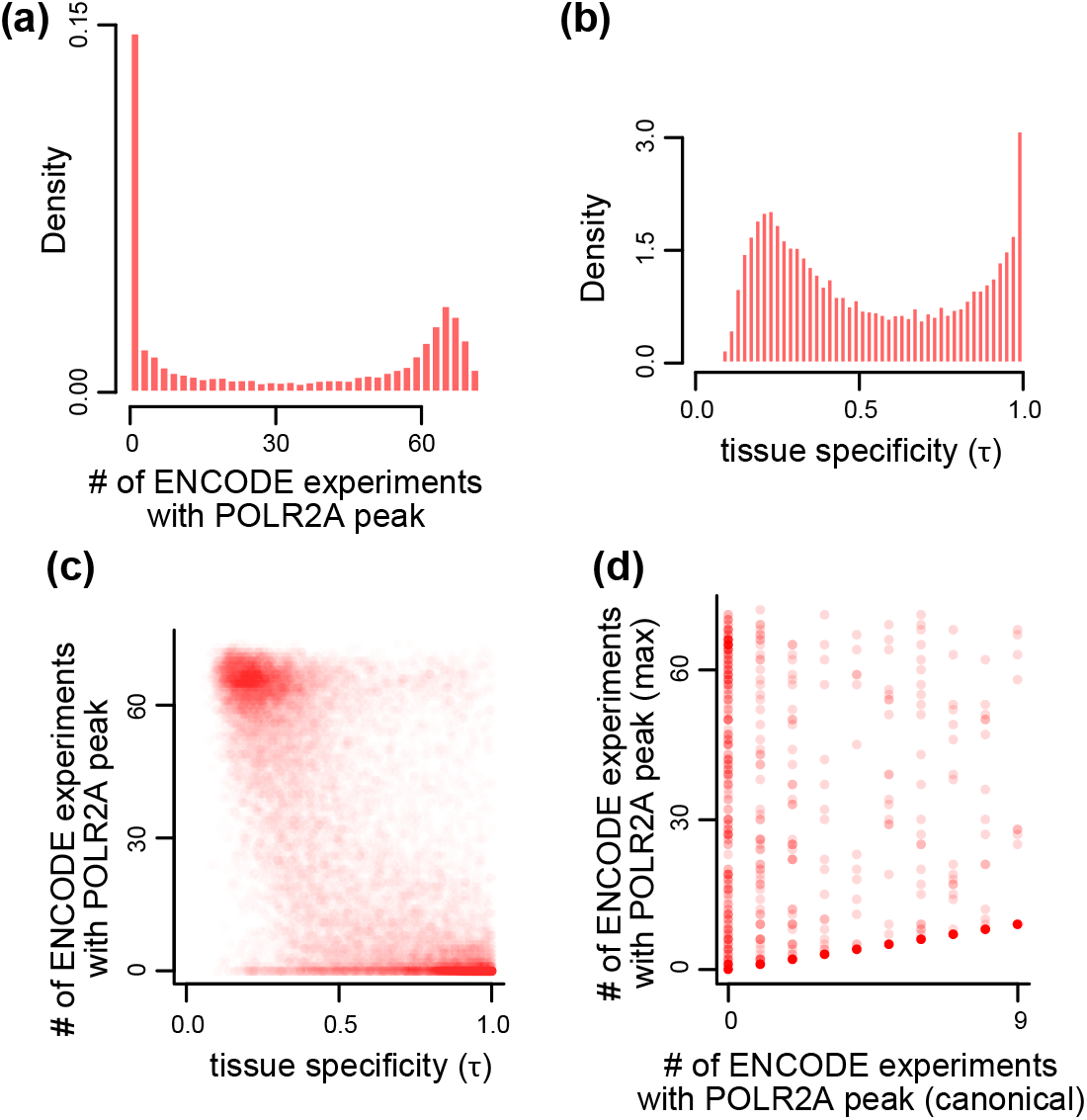
Assessing the relationship between tissue specificity of gene expression and POLR2A binding at the canonical promoter. **(a)** The distribution of the number of ENCODE ChIP-seq experiments showing POLR2A peaks, for all canonical promoters (4 kb regions centered around the TSS) in Ensembl (hg19 assembly). **(b)** The distribution of *τ* computed using gene-level expression quantifications from GTEx. **(c)** Scatterplot of *τ* against the number of ENCODE ChIP-seq experiments showing POLR2A peaks at the canonical promoter. Each point corresponds to a gene-promoter pair. **(d)** Scatterplot of the number of ENCODE ChIP-seq experiments showing POLR2A peaks at the canonical (x axis) promoter versus the corresponding number at the promoter with the greatest number of detected peaks (out of all the alternative promoters of a gene; y axis). Each point corresponds to a promoter pair for a single gene; shown are only genes that are broadly expressed (*τ* < 0.6) but whose canonical promoter shows POLR2A binding in less than 10 ENCODE experiments.

**Supplemental Figure S3.**
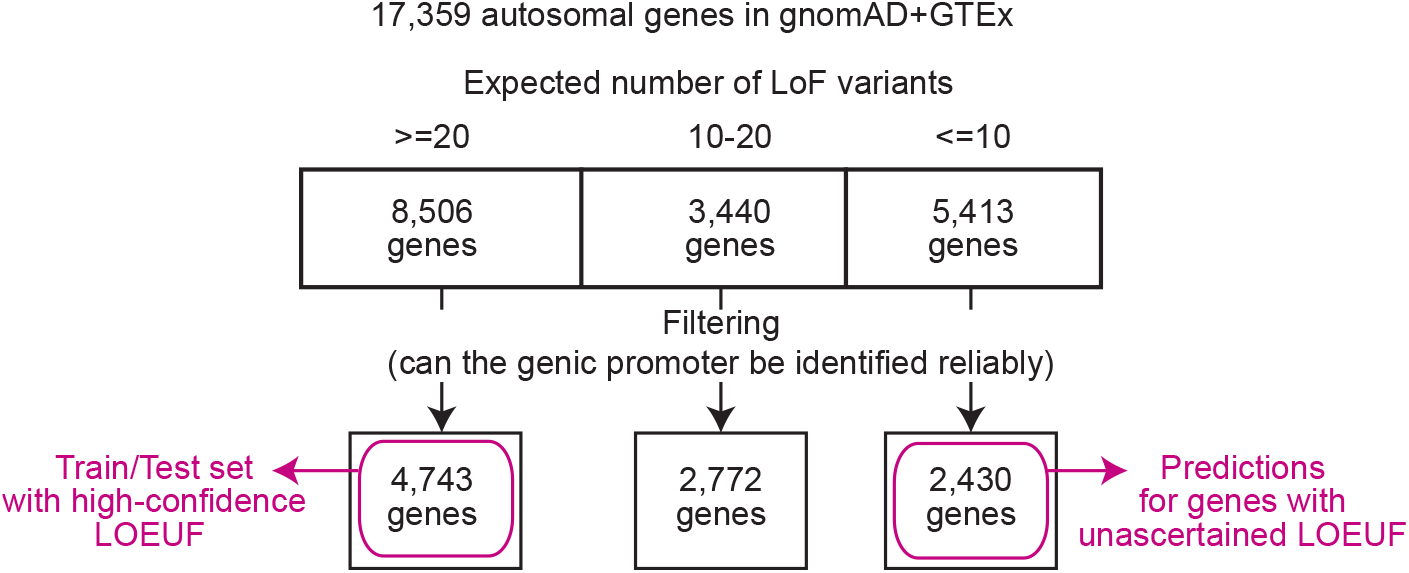
Partitioning genes according to the reliability of their LOEUF estimates and promoter annotation. Schematic illustrating our approach (see Methods for details). We start with 17,359 genes that: a) are present in both GTEx and gnomAD, b) reside in autosomes, and c) their promoters are not subtelomeric. We then filter these according to whether they have reliable promoter annotations, and in cases of pairs of genes with overlapping promoters we only keep one pair. This gives us the set of high-confidence genes that we use to establish the relationship between CpG density and LOEUF and to train predLoF-CpG, and the set of unascertained genes to which we apply predLoF-CpG.

**Supplemental Figure S4.**
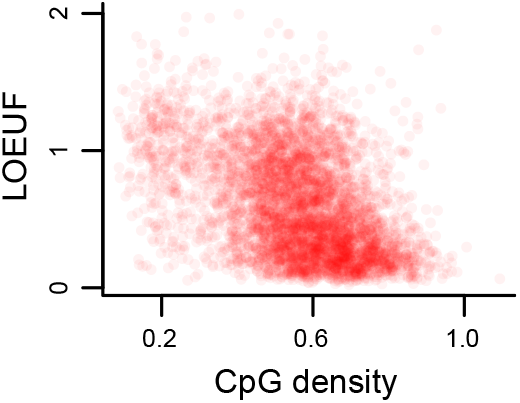
Scatterplot of promoter CpG density against downstream gene LOEUF. Each point corresponds to a promoter-gene pair.

**Supplemental Figure S5.**
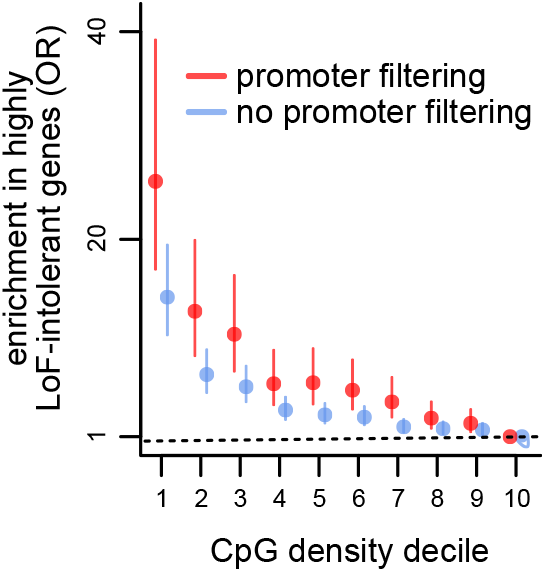
The effect of filtering for high-confidence promoter annotations on the relationship between CpG density and LOEUF. Like Figure 1b, but shown both for the 4,859 genes with high-confidence promoter annotations (red), and for 6,656 genes with canonical (based on GENCODE) promoter annotations and at least 20 expected LoF variants, without further promoter filtering (blue).

**Supplemental Figure S6.**
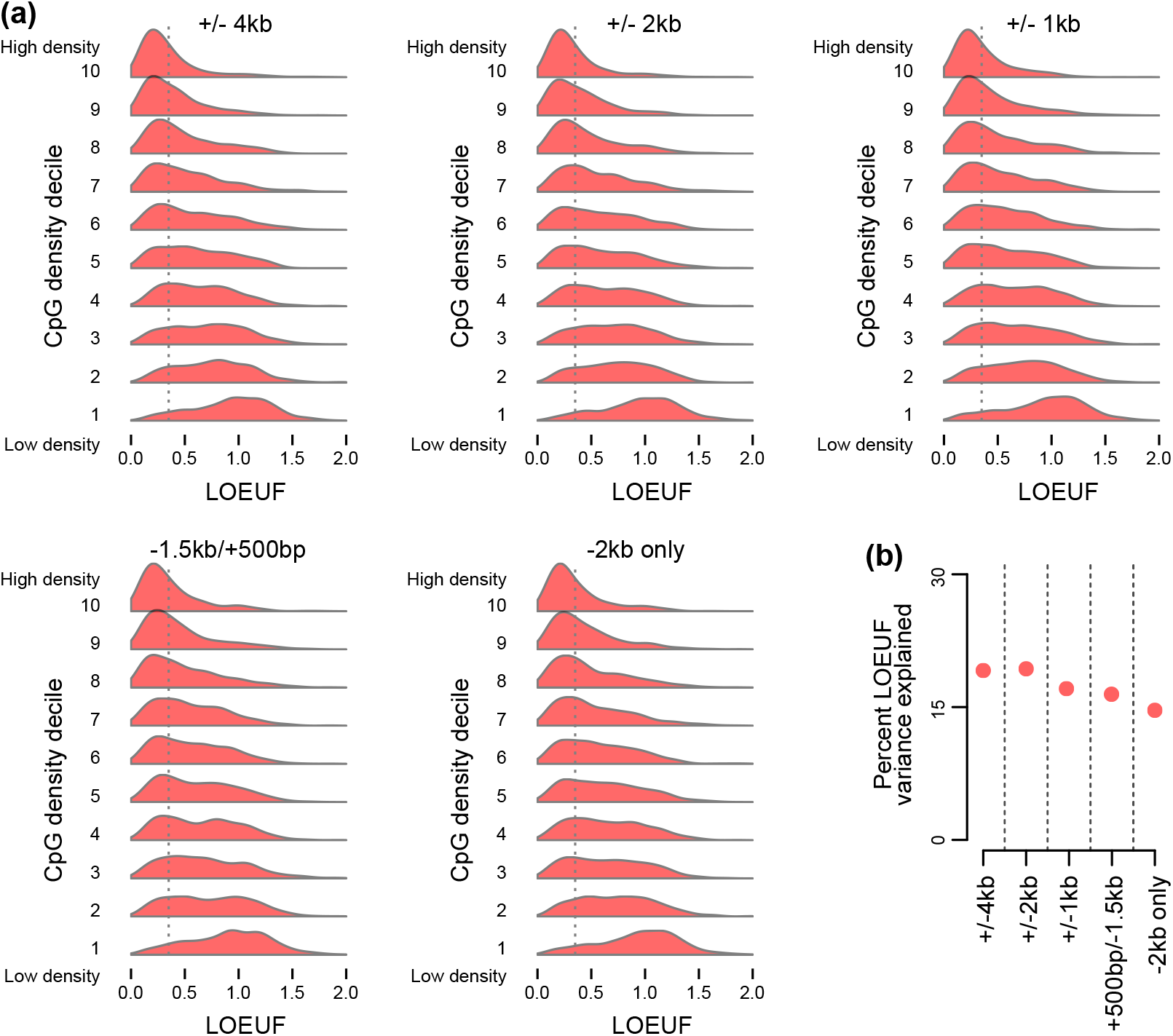
The impact of the size of promoter definition on the relationship between CpG density and LOEUF. **(a)** Like Figure 1a, but with different choices of the interval around the transcription start site that is defined as the promoter. **(b)** The percentage of LOEUF variance (adjusted *r*^2^) that is explained by promoter CpG density, for each of the promoter definitions in (a).

**Supplemental Figure S7.**
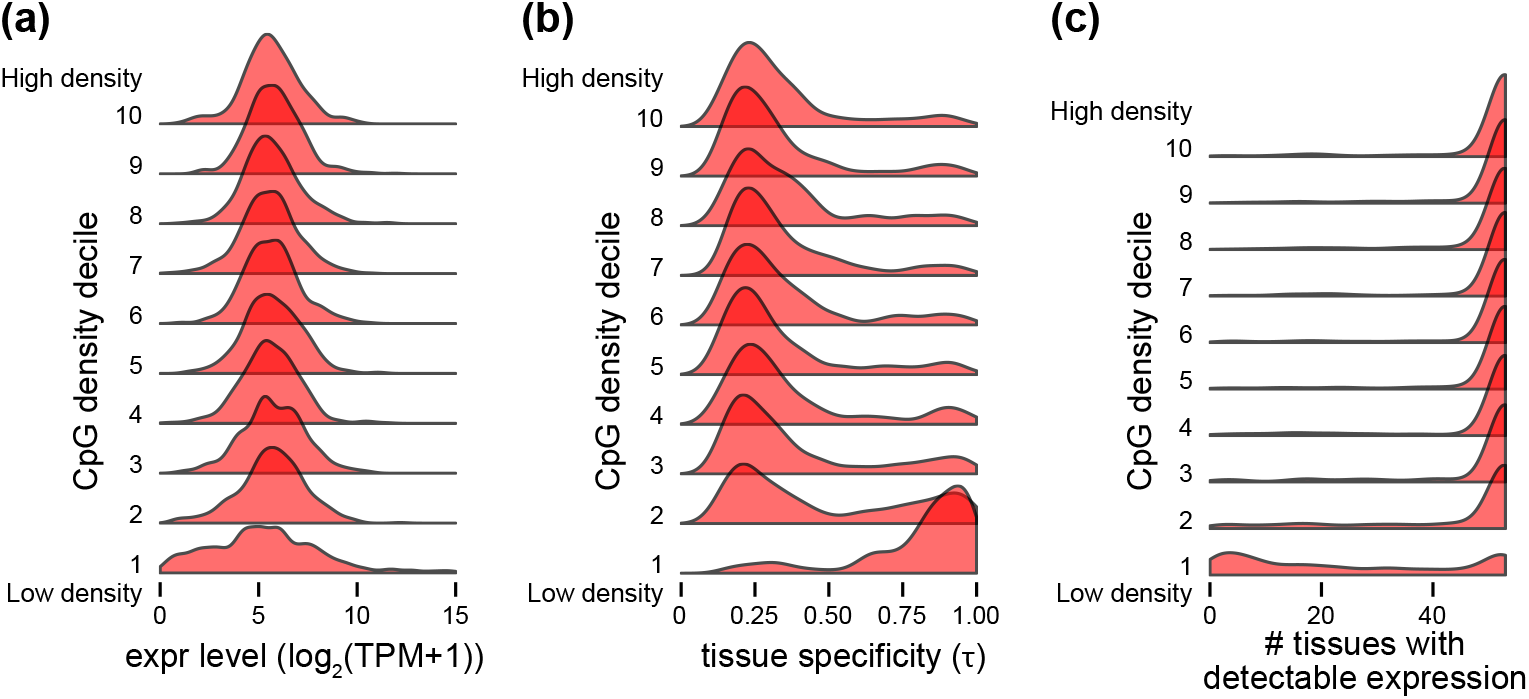
Distributions of downstream gene expression level and tissue specificity across promoter CpG density deciles. Both expression level and *τ* were computed from the GTEx dataset (see Methods). In all three figures, CpG density deciles are labeled 1-10, with 1 the most CpG-poor decile and 10 the most CpG-rich. In **(c)**, detectable expression in a given tissue is defined as median TPM > 0.3.

**Supplemental Figure S8.**
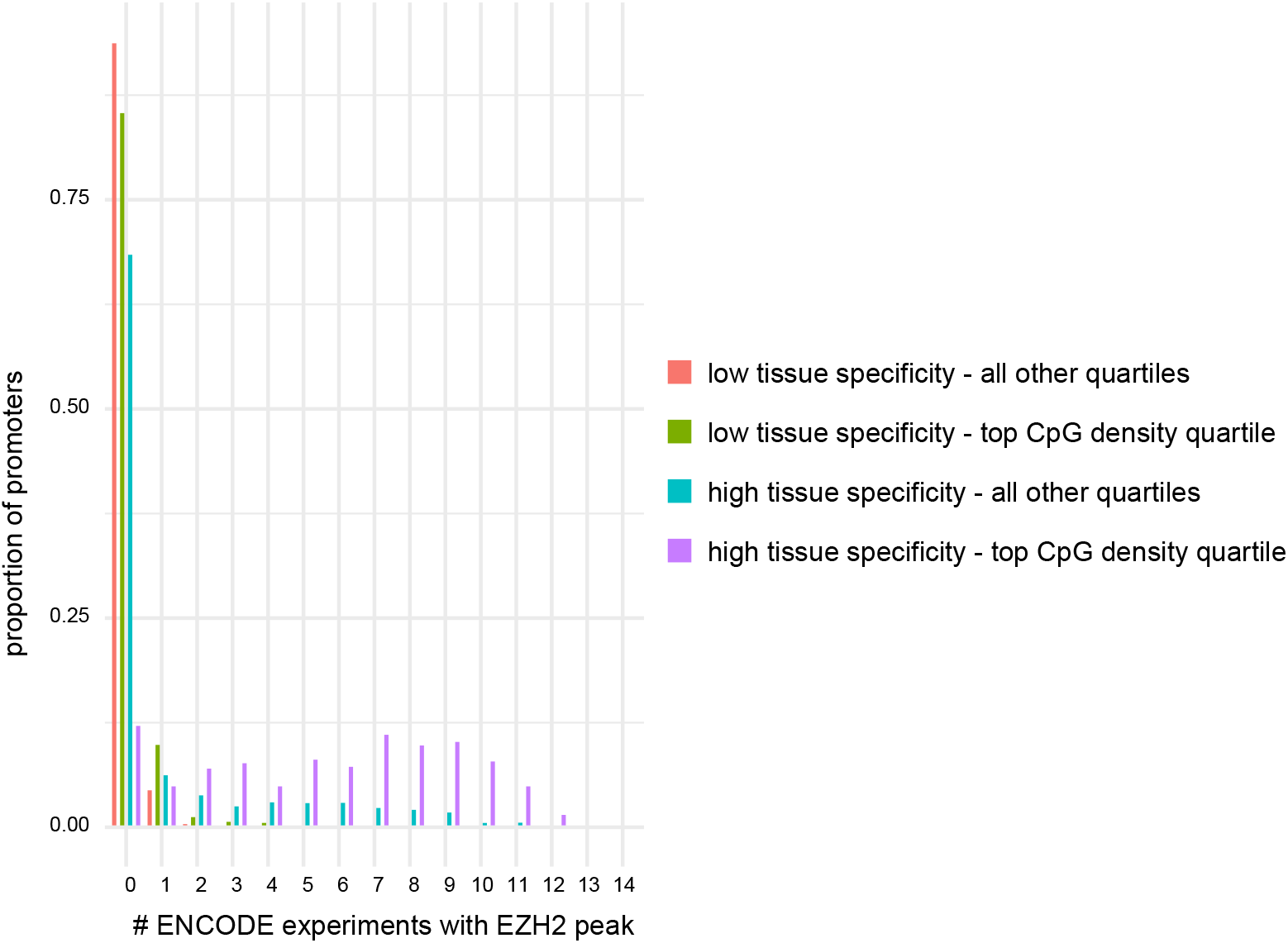
The proportion of promoters with EZH2 peaks in 1-14 ENCODE experiments, stratified based on their CpG density and downstream gene tissue specificity. Tissue speciicity was quantified from the GTEx dataset using *τ* (Methods). Low tissue specificity corresponds to *τ* < 0.6 and high tissue specificity corresponds to *τ* > 0.6.

**Supplemental Figure S9.**
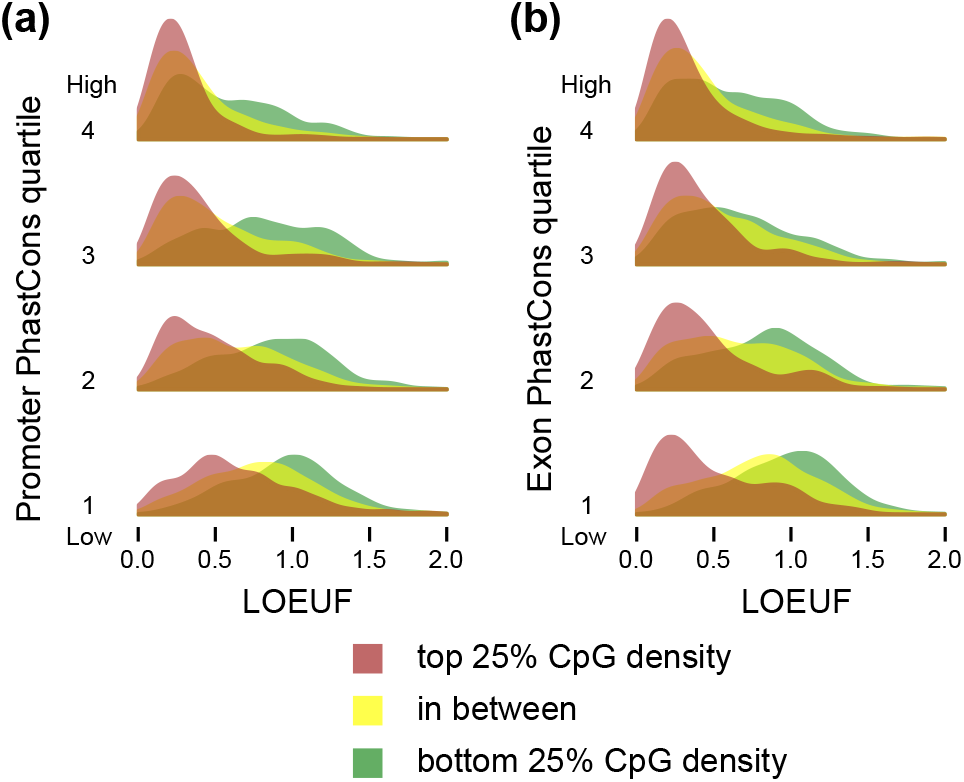
The relationship between promoter CpG density and loss-of-function intolerance conditional on promoter and exonic across-species conservation. **(a)** The distribution of LOEUF, stratified by promoter CpG density, in each quartile of promoter PhastCons score (Methods). **(b)** The distribution of LOEUF, stratified by promoter CpG density, in each quartile of exonic PhastCons (Methods). For both **(a)** and **(b)** quartiles are labeled from 1-4, with 1 being the least and 4 the most conserved, respectively.

**Supplemental Figure S10.**
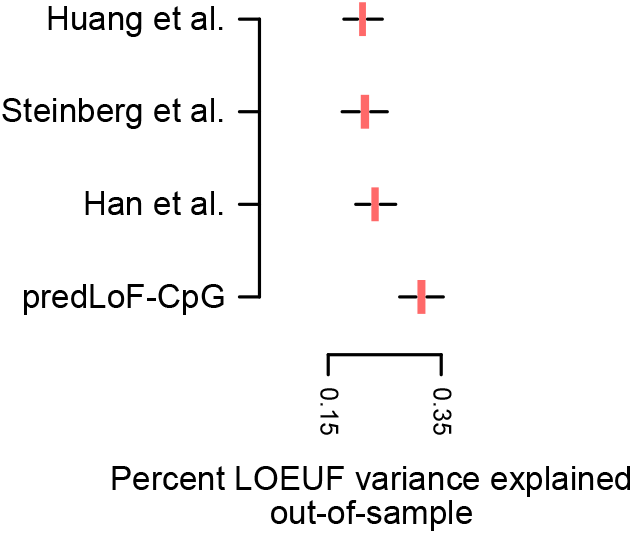
The percentage of out-of-sample LOEUF variance explained by the different predictors of LoF-intolerance. Each boxplots corresponds to a LoF-intolerance predictor as shown on the x-axis, and shows the sampling distribution of the adjusted *r*^2^ after regressing the LOEUF of genes in the test set on the corresponding predictor. We performed 1,000 random train/test splits. For predLoF-CpG, the regression was performed on the prediction probably of high LoF-intolerance.

**Supplemental Figure S11.**
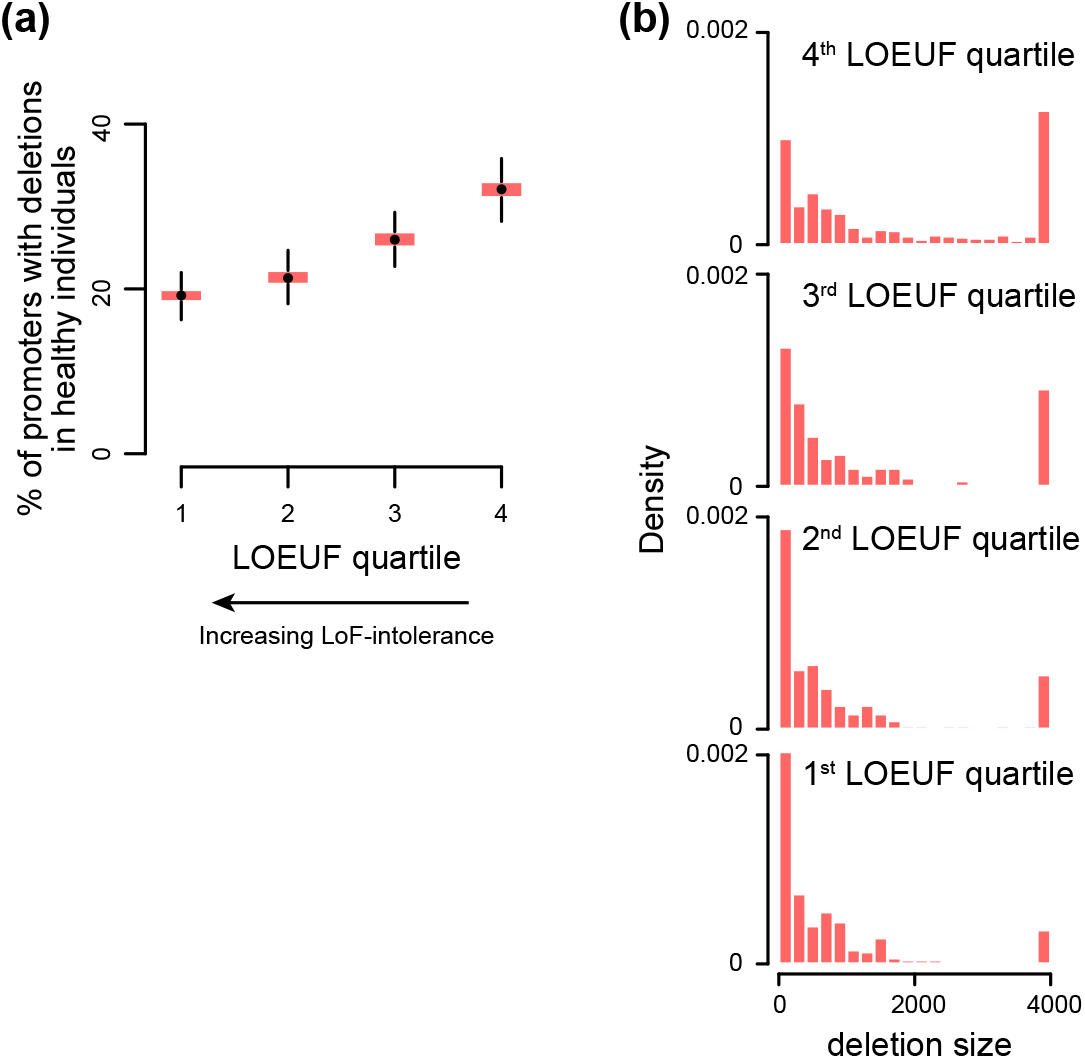
The relationship between promoter deletions seen in healthy individuals and downstream gene loss-of-function intolerance. **(a)** The proportion of promoters harboring deletions across different strata of downstream gene loss-of-function intolerance. For each stratum, the distribution is obtained via the bootstrap. **(b)** The distribution of the size of deletions harbored by promoters across different strata of downstream gene loss-of-function intolerance.

**Supplemental Figure S12.**
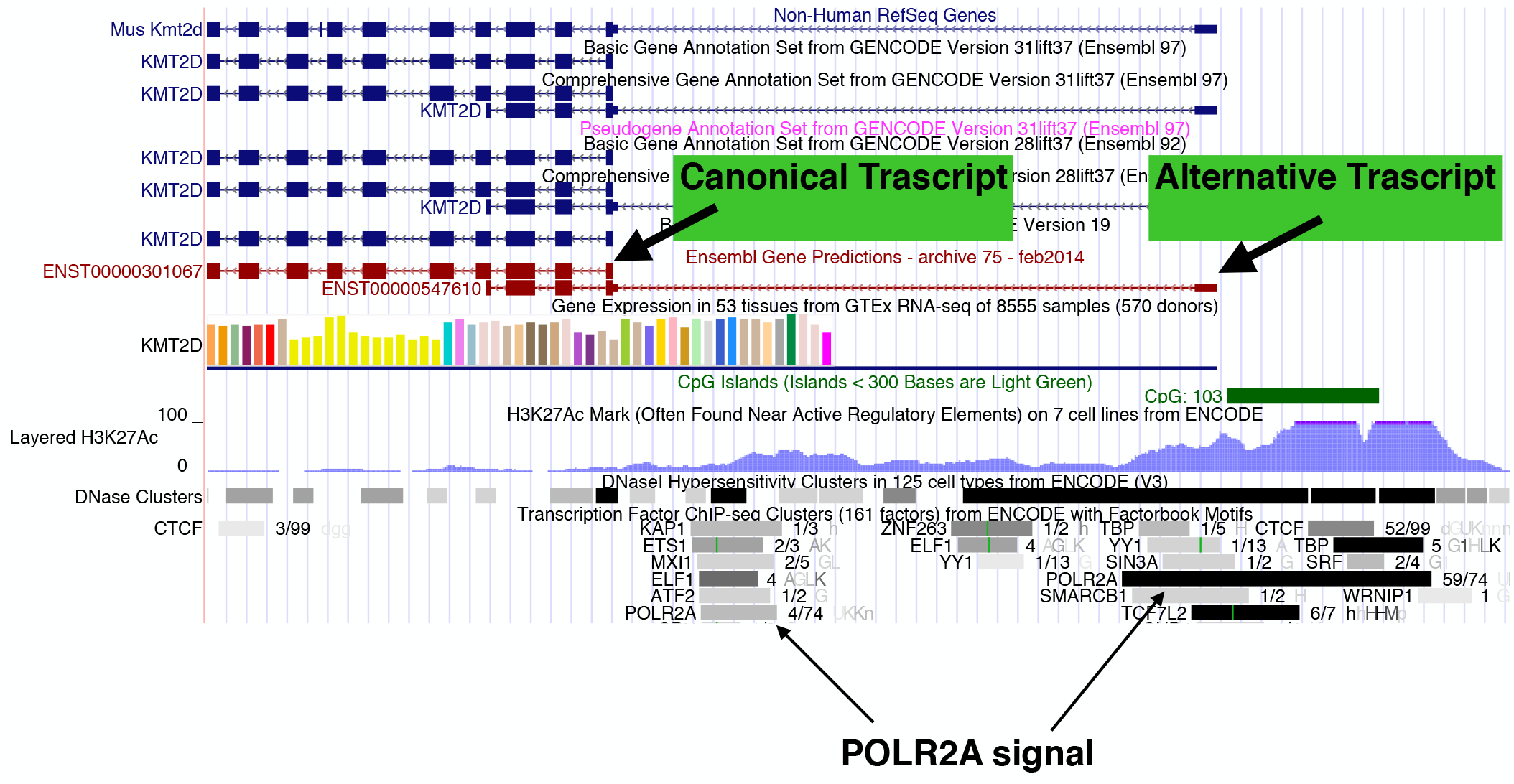
UCSC genome browser screenshot of a 10kb region containing the transcriptional start sites of the canonical and one alternative *KMT2D* trasctipts. The precise coordinates are chr12:49,446,107-49,456,107. The sequence of the canonical transcript extends beyond the 10kb region shown.

## Notes

#### Summary of Updates

Amongst the revisions are - a sensitivity analysis of promoter definition - consideration of genes with number of expected LoF variants between 10 and 20. - a small error was fixed where our set of high-confidence genes inadvertently contained a few with bidirectional promoters - new supplemental table containing all reliable promoter definitions.

